# Medicament identity rather than total loading governs the morphology of electrospun poly(vinylpyrrolidone) nanofibers for regenerative endodontics: a machine learning analysis of a failure-inclusive dataset

**DOI:** 10.64898/2026.09.02.748975

**Authors:** Nura Brimo, Büşra Uysal, Dilek Çökeliler Serdaroğlu

**Affiliations:** Department of Biomedical Engineering, Faculty of Engineering, Başkent University, Bağlıca Campus, 06790 Ankara, Türkiye; Biomedical Engineering, Duke University, Durham, NC 27708, United States; Department of Endodontics, Faculty of Dentistry, Ordu University, 52200 Altınordu/Ordu, Türkiye

**Keywords:** electrospinning, nanofibers, regenerative endodontics, machine learning, active learning, drug delivery

## Abstract

Electrospun fibers loaded with antibiotics or calcium hydroxide are being developed as intracanal carriers for regenerative endodontics, where the dose must stay low enough to spare the stem cells that repopulate the canal. Formulation development sweeps the medicament concentration while holding the polymer and machine settings fixed. We asked whether that sweep targets the right variable. We assembled ENDOSPIN-29, a dataset of 29 poly(vinylpyrrolidone) formulations produced under a single process backbone and loaded with metronidazole, ciprofloxacin, minocycline or calcium hydroxide, alone and in combination, retaining the five that produced no submicron fibers. Across nine regression models, those given per-medicament composition predicted fiber diameter far better than the same models given only total loading. The best reached a leave-one-out coefficient of determination of 0.84 and a median relative error of 18%, whereas every loading-only model performed at or below a mean baseline. Uniformity and distribution span behaved likewise; asymmetry was unpredictable. Dose response ran in opposite directions for different actives: ciprofloxacin thinned fibers monotonically from 406 to 257 nm, while metronidazole thickened them and destroyed fiber formation above 10% w/w. Holding out an entire medicament class removed the advantage, bounding the method to interpolation within a known drug panel.

## Introduction

Pulp necrosis in an immature permanent tooth arrests root development and leaves a thin dentinal wall and an open apex. Regenerative endodontic procedures aim to disinfect such a canal and allow vital tissue to re-form inside it [1,2]. The disinfection step creates a conflict. Triple antibiotic paste, a mixture of metronidazole, ciprofloxacin and minocycline, clears the canal at clinically used concentrations, but those same concentrations kill the stem cells of the apical papilla on which the procedure depends [3,4]. Minocycline also stains dentin [5]. Calcium hydroxide is gentler, but its antibacterial reach inside the canal is limited and removing it from the canal wall before the next treatment stage is difficult.

Electrospun fibers offer a route around this conflict by delivering the same actives from a nanofibrous mat at doses one or two orders of magnitude below a paste [6,7]. Scaffolds carrying ciprofloxacin suppress *Enterococcus faecalis* biofilms without the cytotoxicity of the corresponding paste [8], triple antibiotic fibers act on dual-species biofilms while preserving cell function [9], and the case for fibrous carriers in dental infection control has been reviewed [10]. Fiber diameter is not incidental here. It sets the specific surface area available for release, the mechanical behavior of the mat, and the size of the interconnected pores that cells encounter.

Getting from a drug to a usable mat is a formulation problem. The polymer, the solvent and the machine settings are chosen early and held constant, and the experimentalist sweeps the medicament concentration, recording micrographs after each run to judge whether fibers formed and how thick they are. The sweep is slow, and it rests on an assumption that is rarely stated: that the concentration axis is the axis that matters, so that knowing how much drug a solution carries tells you roughly what will come off the collector.

Machine learning has been applied to electrospinning for about a decade, almost always to one task. Models predict fiber diameter from process variables such as voltage, flow rate, tip-to-collector distance and polymer concentration, trained on data mined from published papers [11,12]. Recent datasets are large; one meta-analysis compiled 68,538 diameter measurements from 1,778 studies across 16 biomedical polymers [11]. Two features of that literature matter here. These are process datasets, and because they pool many polymers and solvents the composition of the loaded cargo is usually reduced to a single concentration figure. They are also assembled by literature mining, which selects for success: runs that produced droplets, beads or nothing rarely appear in a paper, so the boundary of the feasible region is largely absent from the training data.

We report an analysis that inverts both features. We took every formulation recorded in two closely related studies from one laboratory group [13,14], held the process backbone fixed, varied only the loaded medicament, and kept the runs that failed. The dataset is small, with 29 formulations and 26 diameter measurements, and we treat that as a constraint on the claims rather than something to work around. We address five questions: whether medicament identity or total amount governs morphology; whether the same answer holds for the uniformity and shape of the diameter distribution; what dose response a model infers for each active; how far such a model generalizes to a new laboratory and to an unseen medicament class; and what a sequential design algorithm would have done with the same design space.

## Results

### Dataset and design space

All 29 formulations share one process backbone: poly(vinylpyrrolidone) (PVP) at 8% w/w in ethanol, electrospun at 0.9 mL h^-1^ through a 0.4 mm nozzle at a 15 cm tip-to-collector distance under a 0–40 kV field at room temperature. Twenty-five formulations come from the first study [13] and four from a second, independently conducted study [14]. Table 1 lists the set.

**Table 1.** The ENDOSPIN-29 dataset. Concentrations are w/w of the total solution; PVP is 8% w/w throughout, the balance ethanol. *AD* is the mean fiber diameter from *n* = 80 measurements, *SD* its standard deviation, *CV* the ratio. Spinnable is 1 when continuous submicron fibers formed. Laboratory A is ref. 13, B is ref. 14.

| ID | Formulation | MET (%) | CIP (%) | MINO (%) | Ca(OH)<sub>2</sub> (%) | Total (%) | $AD$ (nm) | $SD$ (nm) | $CV$ | Spin. |
| --- | --- | --- | --- | --- | --- | --- | --- | --- | --- | --- |
| N01 | PVP | 0 | 0 | 0 | 0 | 0 | 1087 | 285 | 0.26 | 1 |
| N02 | MET 2.5 | 2.5 | 0 | 0 | 0 | 2.5 | 593 | 105 | 0.18 | 1 |
| N03 | MET 5 | 5 | 0 | 0 | 0 | 5 | 718 | 148 | 0.21 | 1 |
| N04 | MET 7.5 | 7.5 | 0 | 0 | 0 | 7.5 | 636 | 121 | 0.19 | 1 |
| N05 | MET 10 | 10 | 0 | 0 | 0 | 10 | 772 | 192 | 0.25 | 1 |
| N06 | MET 20 | 20 | 0 | 0 | 0 | 20 | 10245 | 3480 | 0.34 | 0 |
| N07 | MET 25 | 25 | 0 | 0 | 0 | 25 | 7797 | 2670 | 0.34 | 0 |
| N08 | CIP 1 | 0 | 1 | 0 | 0 | 1 | 406 | 116 | 0.29 | 1 |
| N09 | CIP 2.5 | 0 | 2.5 | 0 | 0 | 2.5 | 369 | 170 | 0.46 | 1 |
| N10 | CIP 5 | 0 | 5 | 0 | 0 | 5 | 337 | 169 | 0.50 | 1 |
| N11 | CIP 7.5 | 0 | 7.5 | 0 | 0 | 7.5 | 257 | 103 | 0.40 | 1 |
| N12 | CIP 20 | 0 | 20 | 0 | 0 | 20 | – | – | – | 0 |
| N13 | CIP 25 | 0 | 25 | 0 | 0 | 25 | – | – | – | 0 |
| N14 | MET+CIP 5 | 5 | 5 | 0 | 0 | 10 | 804 | 255 | 0.32 | 1 |
| N15 | MET+CIP 7 | 7 | 7 | 0 | 0 | 14 | 744 | 272 | 0.37 | 1 |
| N16 | MET+CIP 9 | 9 | 9 | 0 | 0 | 18 | 642 | 185 | 0.29 | 1 |
| N17 | MET+CIP 11 | 11 | 11 | 0 | 0 | 22 | 690 | 208 | 0.30 | 1 |
| N18 | Triple 5 | 5 | 5 | 5 | 0 | 15 | 1657 | 988 | 0.60 | 1 |
| N19 | Triple 7 | 7 | 7 | 7 | 0 | 21 | 1873 | 985 | 0.53 | 1 |
| N20 | Triple 9 | 9 | 9 | 9 | 0 | 27 | 942 | 682 | 0.72 | 1 |
| N21 | Triple 11 | 11 | 11 | 11 | 0 | 33 | 591 | 465 | 0.79 | 1 |
| N22 | Ca(OH)<sub>2</sub> 2 | 0 | 0 | 0 | 2 | 2 | 545 | 177 | 0.32 | 1 |
| N23 | Ca(OH)<sub>2</sub> 3 | 0 | 0 | 0 | 3 | 3 | 642 | 192 | 0.30 | 1 |
| N24 | Ca(OH)<sub>2</sub> 4 | 0 | 0 | 0 | 4 | 4 | 683 | 244 | 0.36 | 1 |
| N25 | Ca(OH)<sub>2</sub> 5 | 0 | 0 | 0 | 5 | 5 | – | – | – | 0 |
| B01 | PVP (lab B) | 0 | 0 | 0 | 0 | 0 | 885 | 158 | 0.18 | 1 |
| B02 | Ca(OH)<sub>2</sub> 5 (B) | 0 | 0 | 0 | 5 | 5 | 1600 | 1084 | 0.68 | 1 |
| B03 | Ca(OH)<sub>2</sub> 10 (B) | 0 | 0 | 0 | 10 | 10 | 870 | 450 | 0.52 | 1 |
| B04 | Ca(OH)<sub>2</sub> 20 (B) | 0 | 0 | 0 | 20 | 20 | 578 | 397 | 0.69 | 1 |

Concentrations are w/w of the total solution, verified against the recorded solvent masses. A solution of 8% PVP with 9% metronidazole (MET) and 9% ciprofloxacin (CIP) leaves 74% ethanol, and the source records 14.8 g of ethanol in a 20 g solution. Every formulation satisfies this convention, so the composition variables and the solvent fraction are determined exactly rather than estimated. Energy dispersive X-ray spectroscopy confirmed medicament incorporation. Pure PVP fibers gave 63.6 atom % carbon, 13.6% nitrogen and 22.8% oxygen; ciprofloxacin-loaded fibers gave 55.5% carbon, 15.2% nitrogen and 27.0% oxygen together with the fluorine signal that only ciprofloxacin can supply; calcium-loaded fibers showed the calcium signal absent from the polymer alone.

Figure 1 shows representative micrographs. The four panels in the upper row carry between 2.5 and 27% total medicament and all produced continuous submicron fibers. The two lower panels carry 20 and 25% MET and produced beaded structures of 10.2 and 7.8 *µ*m. Total loading does not separate these groups: the triple antibiotic formulation in panel (d) carries far more medicament than either failure and still spins.

**Figure 1.**
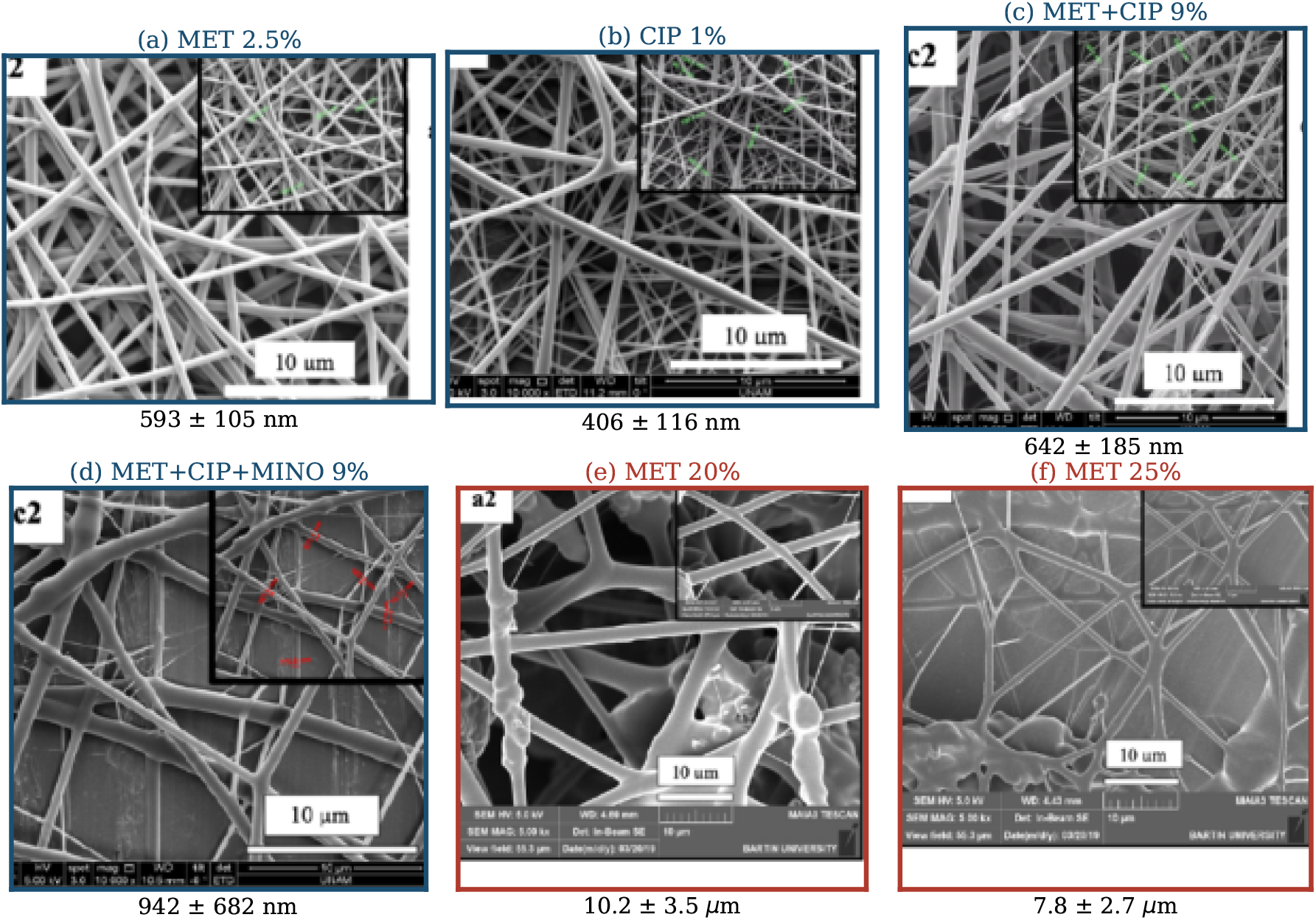
Representative scanning electron micrographs of medicament-loaded PVP fibers, all electrospun under identical conditions. (**a**) 2.5% MET, (**b**) 1% CIP, (**c**) 9% MET with 9% CIP, (**d**) 9% MET with 9% CIP and 9% minocycline, (**e**) 20% MET, (**f**) 25% MET. Panels **a**–**d**, outlined in blue, produced continuous submicron fibers; panels **e** and **f**, outlined in red, produced micron-scale beaded structures and are labeled not spinnable. Values below each panel give the mean fiber diameter and standard deviation from *n* = 80 measurements.

Figure 2a plots the design space. Diameter does not fall monotonically with loading, and the failure boundary is not a loading threshold. Ciprofloxacin drives diameters to 257 nm at 7.5% w/w while the triple antibiotic mixture pushes them above 1.8 *µ*m at 7% of each component. Figure 2b shows the accompanying trade-off: the formulations carrying the largest payload also produced the broadest distributions, with coefficients of variation above 0.7 for the triple antibiotic series.

**Figure 2.**
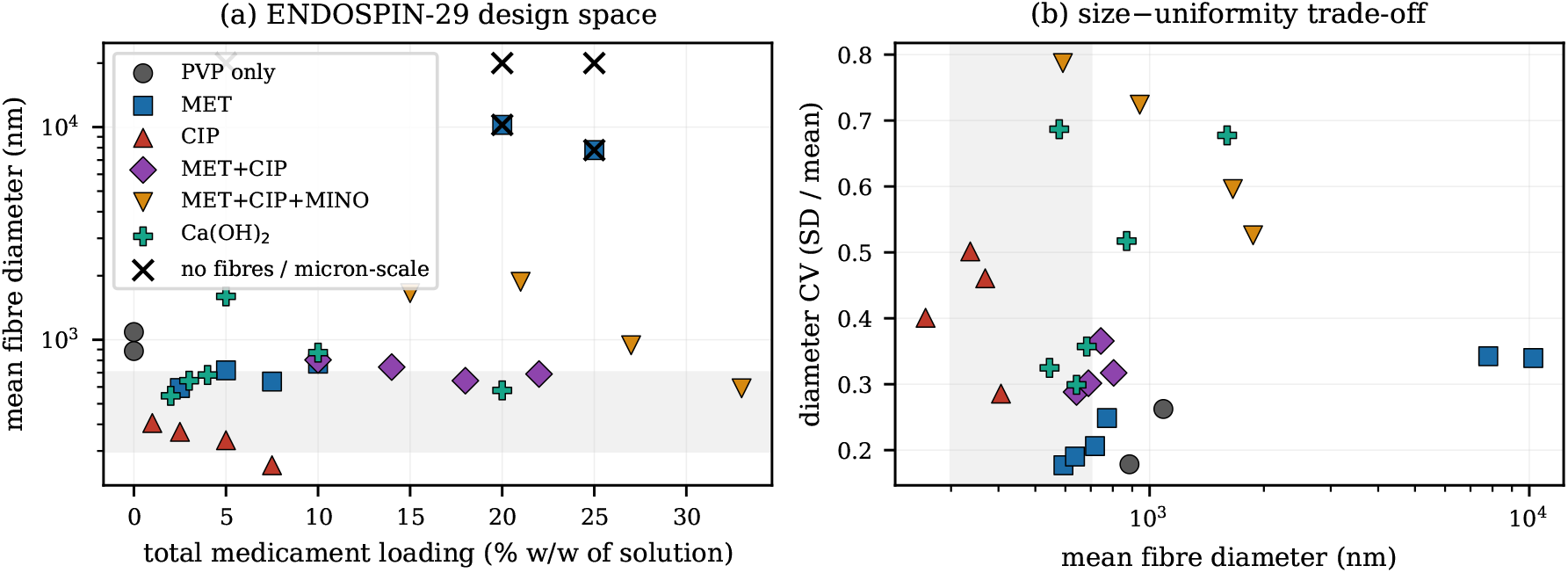
The ENDOSPIN-29 design space. (**a**) Mean fiber diameter against total medicament loading. Crosses mark formulations that produced no submicron fibers; those without a measurable diameter are drawn at the top of the axis. (**b**) Mean diameter against the coefficient of variation. Shaded bands mark the 300–700 nm window used in the design objective.

### Dose response runs in opposite directions for different actives

Within the ciprofloxacin series the diameter falls monotonically with dose, from 406 nm at 1% to 369 nm at 2.5%, 337 nm at 5% and 257 nm at 7.5%, giving a Spearman coefficient of *−*1.00 with no reversals. Within the metronidazole series the diameter rises overall, with a coefficient of +0.89, but reverses direction three times before the structure collapses above 10%. The calcium hydroxide series is not monotone (+0.37) and is confounded by the two contributing laboratories. The double and triple antibiotic series both fall overall (*−*0.80 each), with one reversal apiece.

### Composition predicts fiber diameter; total loading does not

We predicted the decimal logarithm of the mean fiber diameter from three feature sets: total loading alone, the four per-medicament weight fractions, and both. With 26 measurements we used leave-one-out cross-validation throughout and compared nine models against a baseline predicting the training-fold mean.

Table 2 gives the feature ablation and Table 3 the model comparison. Every model given only total loading performed at or below baseline; the Gaussian process reached a coefficient of determination of *−*0.24, worse than ignoring the input entirely. Given per-medicament composition, gradient boosting reached 0.84 with a median relative error of 18%, and the Gaussian process 0.80 with a median relative error of 17%. Six of nine models beat the baseline once composition was available; none did without it. Bootstrap resampling over 10^4^ replicates puts the reduction in mean absolute error for the Gaussian process at 0.155 (95% confidence interval 0.058 to 0.264), and a Wilcoxon signed-rank test on paired absolute errors gives *P* = 0.002. Figure 3 shows the held-out predictions and the ablation.

**Figure 3.**
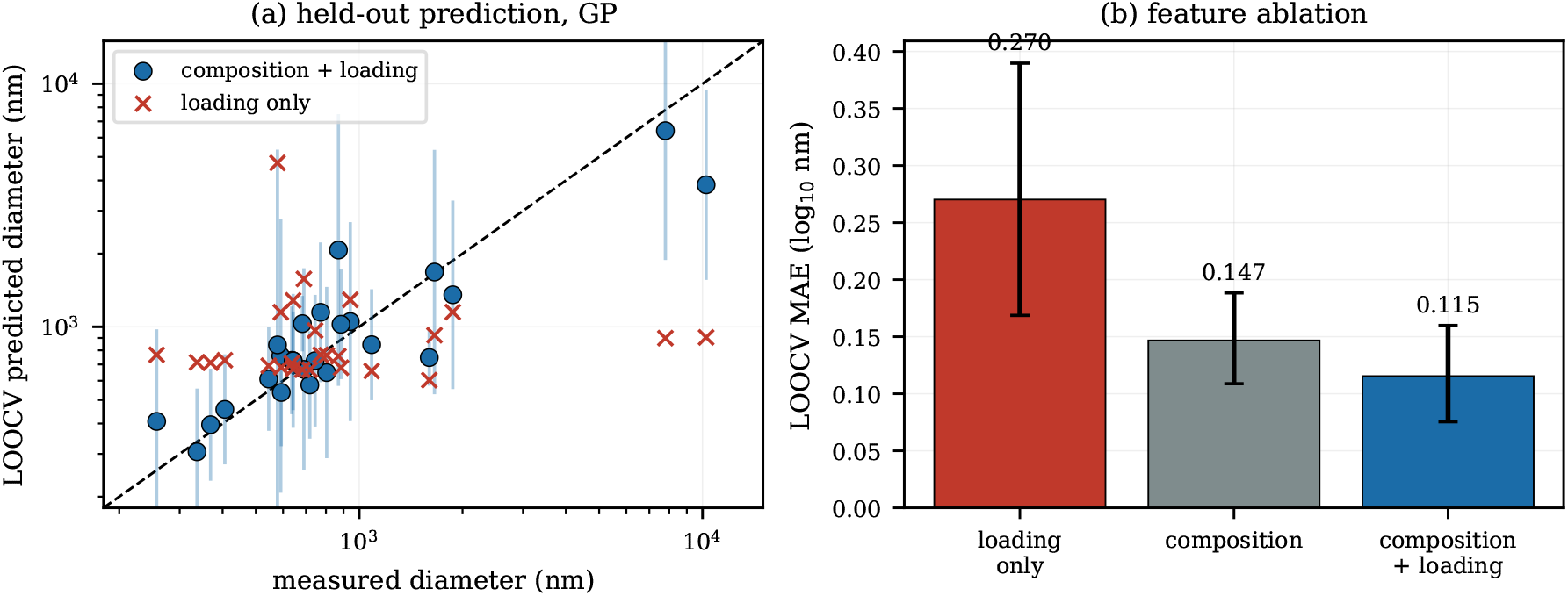
Held-out prediction of fiber diameter. (**a**) Leave-one-out predictions against measured values on logarithmic axes; vertical bars give the *±*1.96*σ* interval of the Gaussian process fitted to composition and loading, and red crosses the predictions of the same model given only total loading. (**b**) Mean absolute error for the three feature sets with bootstrap 95% confidence intervals.

**Table 2.** Feature ablation for the diameter target under leave-one-out cross-validation, *n* = 26. MAE is in decimal logarithmic units. The Spearman coefficient of the mean baseline is *−*1 by construction.

| Model | Features | MAE | $R^2$ | Spearman | Median rel. err. (%) |
| --- | --- | --- | --- | --- | --- |
| Mean baseline | – | 0.248 | $-0.08$ | $-1.00$ | 41.6 |
| Ridge | loading only | 0.253 | $-0.03$ | 0.22 | 46.1 |
| Random forest | loading only | 0.326 | $-0.58$ | 0.16 | 56.5 |
| Gaussian process | loading only | 0.270 | $-0.24$ | 0.08 | 39.0 |
| Ridge | composition | 0.198 | 0.47 | 0.36 | 36.0 |
| Random forest | composition | 0.153 | 0.69 | 0.54 | 33.7 |
| Gaussian process | composition | 0.147 | 0.74 | 0.55 | 28.7 |
| Random forest | composition + loading | 0.148 | 0.68 | 0.63 | 26.3 |
| Gaussian process | composition + loading | **0.115** | **0.80** | **0.81** | **16.8** |

**Table 3.** Model comparison on the diameter target with composition and loading features, leave-one-out cross-validation, *n* = 26, ordered by mean absolute error.

| Model | MAE | $R^2$ | Spearman | Median rel. err. (%) |
| --- | --- | --- | --- | --- |
| Gradient boosting | **0.106** | **0.84** | 0.79 | 18.0 |
| Extra trees | 0.115 | 0.78 | 0.75 | 18.5 |
| Gaussian process (Matérn 5/2) | 0.115 | 0.80 | 0.81 | **16.8** |
| Gaussian process (squared exp.) | 0.128 | 0.76 | 0.77 | 17.9 |
| Support vector regression | 0.134 | 0.68 | 0.72 | 24.4 |
| Random forest | 0.148 | 0.68 | 0.63 | 26.3 |
| Nearest neighbors ($k = 3$) | 0.180 | 0.43 | 0.58 | 26.6 |
| Ridge | 0.204 | 0.43 | 0.35 | 42.0 |
| Lasso | 0.205 | 0.45 | 0.33 | 43.0 |
| Mean baseline | 0.248 | -0.08 | -1.00 | 41.6 |

The pattern repeats for spinnability. Table 4 reports leave-one-out classification of the binary label. A random forest on composition reached an area under the receiver operating characteristic curve of 0.83 against 0.69 for loading alone, with balanced accuracy behaving similarly. The dataset holds 5 negatives against 24 positives, so these estimates are coarse.

**Table 4.** Leave-one-out classification of spinnability, *n* = 29 with 24 positive and 5 negative cases. AUC is the area under the receiver operating characteristic curve.

| Model | Features | AUC | Balanced accuracy |
| --- | --- | --- | --- |
| Majority baseline | – | 0.500 | 0.500 |
| Logistic regression | loading only | 0.683 | 0.754 |
| Random forest | loading only | 0.688 | 0.796 |
| Logistic regression | composition | 0.717 | 0.817 |
| Logistic regression | composition + loading | 0.767 | 0.838 |
| Random forest | composition + loading | 0.750 | 0.658 |
| Random forest | composition | **0.829** | **0.858** |

### Uniformity and span follow the same pattern; asymmetry does not

We repeated the analysis for three further targets: the coefficient of variation, the relative span (maximum minus minimum, divided by the mean), and an asymmetry index positive when the distribution has a long upper tail (Table 5). Uniformity is predictable from composition, with a coefficient of determination of 0.54 against *−*0.09 for loading alone and *−*0.08 for baseline. Relative span reaches 0.66 against 0.25 for loading. Asymmetry is not predictable from either feature set, at *−*0.07 and *−*0.25 respectively, both at or below baseline.

**Table 5.** Prediction of three further morphological targets by the Gaussian process under leave-one-out cross-validation.

| Target | Features | $n$ | MAE | $R^2$ |
| --- | --- | --- | --- | --- |
| Coefficient of variation | mean baseline | 26 | 0.151 | -0.08 |
|  | loading only | 26 | 0.152 | -0.09 |
|  | composition + loading | 26 | **0.094** | **0.54** |
| Relative span | mean baseline | 22 | 0.615 | -0.10 |
|  | loading only | 22 | 0.514 | 0.25 |
|  | composition + loading | 22 | **0.301** | **0.66** |
| Distribution asymmetry | mean baseline | 22 | 0.159 | -0.10 |
|  | loading only | 22 | 0.155 | -0.25 |
|  | composition + loading | 22 | 0.152 | -0.07 |

### Inferred dose response, data efficiency and class extrapolation

Figure 5a shows the dose response the fitted Gaussian process infers for each active with the others held at their dataset values. Ciprofloxacin drives the predicted diameter from about 1450 nm at zero to a minimum near 600 nm around 10–12% w/w. Metronidazole does the reverse, rising from about 720 nm to above 4000 nm by 20%, which is where the measured structures failed. Calcium hydroxide falls gently and monotonically. Minocycline shows a shallow decline across the narrow range tested.

The learning curve in Figure 5b shows mean absolute error falling from 0.234 at 6 training formulations to 0.119 at 24, with no plateau.

Figure 5c reports a harder test. Holding out an entire medicament class rather than a single formulation removes the advantage. The overall coefficient of determination for the Gaussian process falls to *−*0.06, and the median relative error on the held-out ciprofloxacin series rises to 91%. Only the double antibiotic and calcium hydroxide classes, both of which retain close relatives in the training set, are predicted better than the leave-one-out error.

### Uncertainty calibration and cross-laboratory transfer

Empirical coverage of the Gaussian process intervals under leave-one-out cross-validation was 0.65 at a nominal 0.50 and 0.85 at a nominal 0.80, conservative in the middle of the distribution. At a nominal 0.95 the empirical coverage was 0.88, with a standard deviation of the standardized residuals of 1.44.

Training on the 22 formulations from the first laboratory and testing on the 4 calcium hydroxide formulations from the second gave a mean absolute error of 0.161, close to the internal cross-validated 0.149. The aggregate hides a disagreement. Both laboratories prepared calcium hydroxide at 5% w/w with the same polymer and solvent. In the first, no fibers formed. In the second, the same nominal composition produced fibers of 1600 *±* 1084 nm.

### Functional consequences of the fiber format

Data from the second study [14] is summarized in Table 6. Against *E. faecalis*, calcium hydroxide alone showed no inhibition at 1000 *µ*g mL^-1^ against either strain tested. The PVP fiber carrier inhibited both strains at 1000 *µ*g mL^-1^, and the calcium hydroxide loaded fiber at 125 *µ*g mL^-1^, an eightfold reduction. None of the preparations acted on *E. coli* or *P. aeruginosa* at the concentrations tested. In an MTT assay on NIH/3T3 fibroblasts over 24 h, no preparation differed significantly from the untreated control at 0.1, 1 or 2 mg mL^-1^ (*P >* 0.05, *n* = 5).

**Table 6.** Functional data for the calcium hydroxide system from ref. 14. Minimum inhibitory concentrations in *µ*g mL^-1^; residual medicament as Kruskal-Wallis mean ranks over 224 canals, lower values indicating less residue. GP denotes a gutta-percha cone.

| *Minimum inhibitory concentration* |  |  |  |
| --- | --- | --- | --- |
| Organism | PVP fiber | Ca(OH)<sub>2</sub>-PVP fiber | Ca(OH)<sub>2</sub> |
| *E. faecalis* ATCC 29212 | 1000 | **125** | >1000 |
| *E. faecalis* ATCC 51299 | 1000 | **125** | >1000 |
| *E. coli* ATCC 25922 | >1000 | >1000 | >1000 |
| *P. aeruginosa* ATCC 27853 | >1000 | >1000 | >1000 |
| *Residual medicament after removal ($n = 56$ per group, $P < 0.001$)* |  |  |  |
| Group | Mean rank | Syringe vs ultrasonic ($P$) | |
| Ca(OH)<sub>2</sub> powder and liquid | 160.8 | <0.001 |  |
| Ca(OH)<sub>2</sub> injectable | 143.0 | <0.001 |  |
| GP with Ca(OH)<sub>2</sub>-PVP fiber | 77.5 | 0.670 |  |
| GP with PVP fiber | **68.6** | 0.009 |  |

The removal study used 224 extracted upper incisors in eight groups. Ranked by residual material, mean ranks were 160.8 for calcium hydroxide powder and liquid, 143.0 for the injectable paste, 77.5 for the fiber-coated cone carrying calcium hydroxide and 68.6 for the polymer fiber-coated cone, with paste-to-fiber differences significant at *P <* 0.001. Passive ultrasonic irrigation removed more residue than syringe irrigation overall (mean ranks 93.8 against 131.2 across 112 canals each). Broken down by group, the advantage of ultrasonic activation was large for both pastes (*P <* 0.001 each), smaller for the polymer fiber group (*P* = 0.009), and absent for the calcium hydroxide loaded fiber group (*P* = 0.670).

### Sequential design over the measured pool

Each formulation was scored by a desirability function combining proximity of the diameter to a 300–700 nm window, uniformity, and payload as a geometric mean, with non-spinnable formulations scoring zero. A Gaussian process seeded with four randomly chosen formulations then selected from the remaining pool by expected improvement or an upper confidence bound, over 200 random seeds. Under the primary objective, expected improvement reached the pool optimum after 7.5*±* 0.2 runs and the upper confidence bound after 7.4 *±* 0.2, against 14.3 *±* 0.5 for random selection (Figure 4). The advantage was 1.7-fold under a payload-first objective and 1.2-fold under a payload-agnostic one, where the optimum is an isolated low-loading formulation.

**Figure 4.**
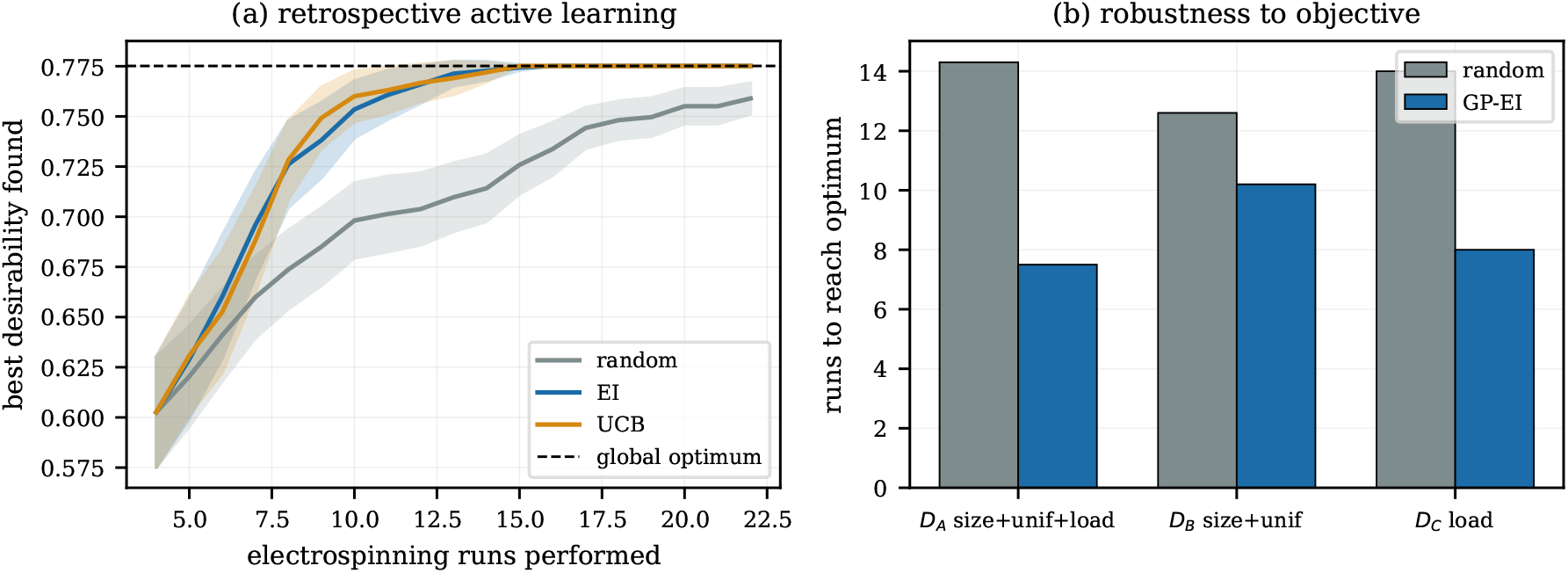
Retrospective sequential design. (**a**) Best desirability found against the number of electrospinning runs performed, averaged over 200 random seeds with 95% confidence bands. (**b**) Runs required to reach the optimum under three objectives.

**Figure 5.**
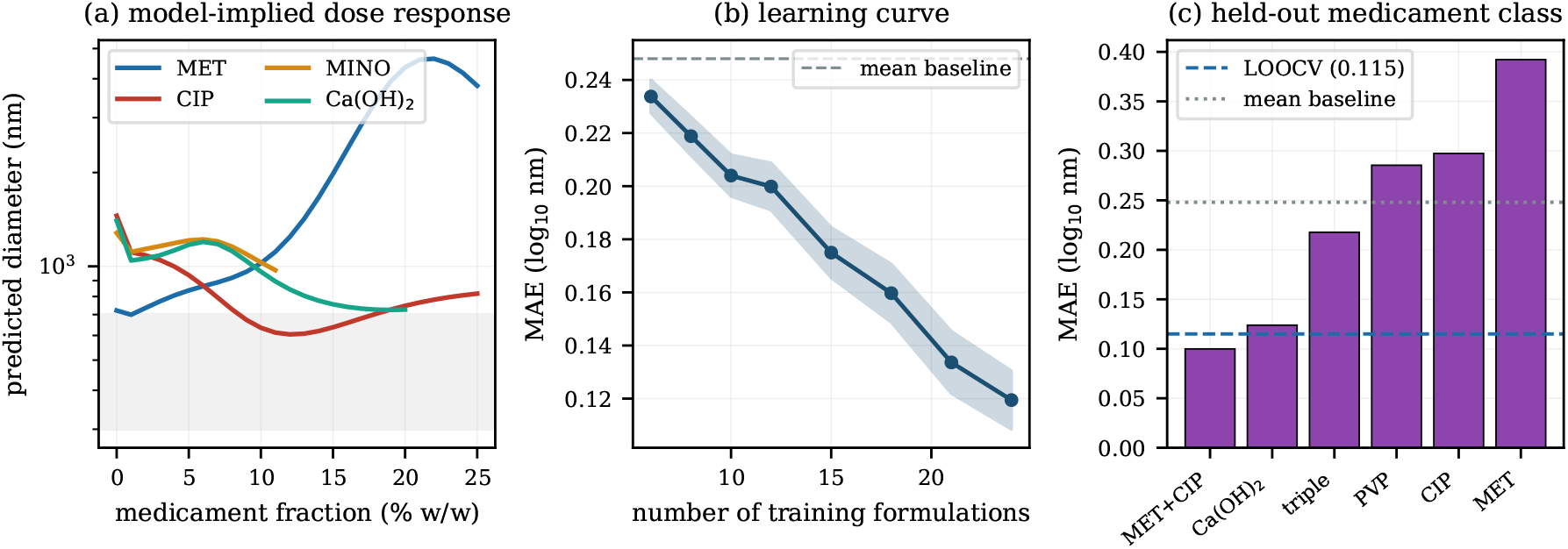
Model behavior and its limits. (**a**) Dose response inferred by the fitted Gaussian process for each active, with the shaded band marking the 300–700 nm window. (**b**) Learning curve, mean absolute error against the number of training formulations, over 200 random splits with a 95% band. (**c**) Mean absolute error when an entire medicament class is held out, compared with the leave-one-out error and the mean baseline.

The source studies independently selected three formulations for downstream work by visual inspection of micrographs: MET with CIP at 9% each, the triple antibiotic at 9% each, and calcium hydroxide at 3%. Under our primary objective these rank 2nd, 9th and 15th of 26. The model’s own first choice, MET with CIP at 11% each, was not carried forward.

## Discussion

The central result is that two medicaments in the same polymer and the same solvent move the fiber diameter in opposite directions, and that a representation summing them discards this. Ciprofloxacin thins the fibers monotonically; metronidazole thickens them and then destroys fiber formation altogether. Every model we tested reproduced this separation when given per-medicament composition, and none recovered it from the total loading.

The mechanistic reading is that ciprofloxacin raises the conductivity of the ethanol solution, increasing the charge density on the jet and the stretching it undergoes before solidification. Metronidazole appears instead to approach its solubility limit, so that above roughly 10% w/w it comes out of solution and disrupts the jet, producing beaded micron-scale structures rather than thinner fibers. Calcium hydroxide is a suspended powder rather than a dissolved solute, which is consistent with the different behavior it showed across the two laboratories. The model recovered these directions from data alone, without mechanistic input.

The practical consequence is direct. A concentration sweep that varies one drug across four levels traverses the axis carrying the least information while holding constant the variable carrying the most. Sampling across actives at fewer levels each would map the same design space in fewer runs, and the sequential design simulation suggests roughly half as many.

The failed runs earn their place in the dataset. Metronidazole at 20% and ciprofloxacin at 20% both fail at identical total loading, for different reasons and with different signatures, while the triple antibiotic at 27% total succeeds. No model trained only on successes can represent that boundary, and no literature-mined dataset contains it. This is our main methodological point against the prevailing practice of assembling electrospinning datasets from published papers [11,12]: the resulting collections are survivorship-biased by construction, however large they become.

The class extrapolation result sharpens rather than undermines the main claim. Composition features work by interpolating within the space of medicaments the model has seen. They confer no ability to predict a drug absent from training, because the model has no descriptor for what a new molecule will do to solution conductivity or solubility. A model built to screen genuinely new actives would need molecular descriptors and training data spanning enough chemistry for those descriptors to be identifiable. Within a defined panel of actives, which is the situation in regenerative endodontics where the same handful of drugs recur, interpolation is the relevant task.

The asymmetry result bounds the claim in a second way. Composition carries information about how wide the diameter distribution will be but not about which side of the mean its tail falls on. Skewness here is plausibly governed by local jet instabilities that formulation variables do not capture, and resolving it would require per-fiber measurements rather than summary statistics.

The two laboratories disagreed at a shared design point, with 5% calcium hydroxide producing no fibers in one and 1600 nm fibers in the other. We cannot resolve this from the available records; ambient humidity, PVP molecular weight grade, calcium hydroxide particle size and dispersion during stirring are all plausible contributors and none was logged. Calcium hydroxide is the one medicament in the set that remains a suspension, making it the formulation most sensitive to exactly these unrecorded variables. The episode cautions against reading a small cross-laboratory error as evidence of generality.

The functional data connects morphology to outcomes that matter clinically. The eightfold reduction in the concentration required for *E. faecalis* inhibition, and the roughly halved residual material after removal, indicate that the fiber format is not merely a tidier way to present the same drug. The absence of any significant difference between syringe and ultrasonic irrigation for the calcium hydroxide fiber group (*P* = 0.670), where both pastes showed large differences, suggests the format is also less dependent on which irrigation protocol is available. These endpoints were measured on a few selected formulations rather than across the design space, so we did not model them.

Several limitations constrain the interpretation. The dataset has 29 formulations, 26 diameters and 5 negatives, so confidence intervals are wide and the classification results rest on few failures. The learning curve indicates the model remains data-limited, making the reported accuracy a lower bound rather than a converged figure. Diameters are recorded as summary statistics rather than the underlying 80 measurements per sample, so the asymmetry index is a coarse proxy. Spinnability labels derive from textual descriptions by a fixed rubric, and a different rubric would move borderline cases. The elemental analyses confirm incorporation but do not quantify loading efficiency, since energy dispersive X-ray spectroscopy is surface-weighted and semi-quantitative on light elements. The sequential design result is retrospective, showing that a Gaussian process would have navigated an already measured pool efficiently, which is necessary but not sufficient for prospective gain, and it treats failed and successful runs as equally costly. Finally, the two source studies share an institution and a supervisor, so the cross-laboratory analysis understates the difficulty of moving a model between independent groups.

One inconsistency in the source material requires recording. The histogram inset for the 9% MET with CIP formulation is annotated 942 *±* 185 nm while the accompanying text gives 642 *±* 185 nm. The plotted distribution spans 0.4–1.2 *µ*m, consistent with the smaller value, and 942 nm is the figure reported for the 9% triple antibiotic sample. We used 642 nm.

The immediate test is prospective. The model ranks MET with CIP at 11% each above the formulations that were carried forward, and the learning curve indicates an additional measurement is worth more to the model now than at any later point. A single confirmatory run would settle whether that ranking holds.

## Methods

### Materials and solution preparation

PVP was used as the carrier polymer at 8% w/w in all formulations, with ethyl alcohol as solvent. Metronidazole, ciprofloxacin, minocycline and calcium hydroxide were used as received. For each formulation the medicament and PVP powders were dissolved together in the stated mass of ethanol at room temperature and stirred on a magnetic stirrer for two nights with the vessel closed to air, then drawn into 2 mL plastic syringes. Concentrations are w/w of the total solution, so the ethanol fraction is 100% minus the polymer and medicament fractions. Single medicament series covered MET at 2.5, 5, 7.5, 10, 20 and 25%; CIP at 1, 2.5, 5, 7.5, 20 and 25%; and calcium hydroxide at 2, 3, 4 and 5%. The double antibiotic series used equal fractions of MET and CIP at 5, 7, 9 and 11% each, and the triple antibiotic series equal fractions of MET, CIP and minocycline at the same four levels. A second laboratory prepared calcium hydroxide at 5, 10 and 20% with the same polymer and solvent [14].

### Electrospinning

A single-nozzle system was used throughout, at a flow rate of 0.9 mL h^-1^, nozzle inner diameter 0.4 mm, and a 15 cm distance between nozzle and grounded collector. The high-voltage supply operated over 0–40 kV with the positive electrode connected to the metallic needle and the ground to the collector, a rectangular metal plate covered with aluminum foil. Collection time was 15 s, at room temperature.

### Characterization

Fiber morphology was examined by scanning electron microscopy on a FEI Quanta 200 FEG ESEM at approximately 12 kV, after coating with an Au/Pd layer of about 5 nm using a Gatan 682 precision etching and coating system. Images were recorded at 1500-fold and 10,000-fold magnification. Mean fiber diameters were obtained from *n* = 80 measurements at different locations using ImageJ, and are reported as mean with standard deviation. Chemical composition was checked by energy dispersive X-ray spectroscopy on a Hitachi SU1510 system at 2000-fold magnification. Full characterization of the antibiotic-loaded series, including X-ray photoelectron spectroscopy of coated gutta-percha cones, has been reported [15].

### Antibacterial, cytotoxicity and removal assays

Minimum inhibitory concentrations were determined against *E. faecalis* ATCC 29212 and ATCC 51299, *E. coli* ATCC 25922 and *P. aeruginosa* ATCC 27853 by broth dilution up to 1000 *µ*g mL^-1^. Cytotoxicity was assessed on NIH/3T3 fibroblasts by MTT assay after 24 h, absorbance read at 540 nm, viability expressed relative to a growth medium control, *n* = 5 per group, compared by one-way analysis of variance. For the removal study, 224 human upper incisors were decoronated to 16 mm, prepared with a Reciproc Blue R40 file, and assigned at random to eight groups of 28 defined by intracanal medicament and removal method. Residual material was compared by Kruskal-Wallis and Mann-Whitney U tests. Full protocols appear in the source work [14]. The extracted teeth were obtained and handled under the ethical approval granted for that study; no living human or animal subjects were involved in the work reported here.

### Dataset construction and labeling

Each row of ENDOSPIN-29 records the four per-medicament weight fractions, the total loading, the number of distinct actives, the laboratory of origin, the mean fiber diameter with standard deviation and, where reported, minimum and maximum diameters, plus a binary spinnability label. The coefficient of variation is the standard deviation divided by the mean; the relative span is the difference between maximum and minimum divided by the mean; the asymmetry index is the difference between upper and lower deviations from the mean divided by the full range.

Spinnability was assigned 1 when the source reports continuous submicron fiber structures, and 0 when it reports no fiber structures or fibers only at micron scale with droplets or beads dominating. This produced five negatives: MET at 20 and 25%, CIP at 20 and 25%, and calcium hydroxide at 5% in the first laboratory. Three lack a reported diameter and contribute only to classification. Labels were taken from explicit statements in the source text rather than from graded morphological adjectives.

### Machine learning

All models were implemented with scikit-learn 1.8 [16]. The regression target was the decimal logarithm of the mean diameter. The Gaussian process used a constant kernel multiplied by a Matérn kernel with *ν* = 5*/*2 plus a white noise term, inputs standardized, target normalized, five optimizer restarts. A second Gaussian process used a squared exponential kernel. Tree ensembles used 500 estimators; gradient boosting used library defaults; ridge and lasso used *α* = 1 and *α* = 0.01; the support vector regressor used a radial basis kernel with *C* = 10 and *ɛ* = 0.05; the nearest neighbor regressor used *k* = 3.

Evaluation used leave-one-out cross-validation. Confidence intervals came from bootstrap resampling of held-out absolute errors over 10^4^ replicates, and feature sets were compared by a Wilcoxon signed-rank test on paired absolute errors. The learning curve drew 200 random training subsets at each size, evaluating on the complement. For class extrapolation, all formulations of one medicament class were removed from training and predicted from the remainder. Classification used logistic regression and a random forest with balanced class weights, scored by area under the receiver operating characteristic curve and balanced accuracy.

For sequential design, desirability was the geometric mean of a size term (1 inside the 300–700 nm window, decaying linearly below and exponentially above), a uniformity term (1 *− CV* clipped to the unit interval) and a payload term (total loading normalized by the dataset maximum), with non-spinnable formulations assigned zero. A Gaussian process seeded with four random formulations selected from the remaining pool by expected improvement or an upper confidence bound with coefficient 2, until the pool optimum was selected or the pool exhausted, averaged over 200 seeds. Two alternative objectives were evaluated identically, one omitting the payload term and one using payload alone subject to a 1000 nm ceiling.

### Use of large language models

A large language model was used to assist with drafting and language editing of this manuscript, and to assist in writing the analysis code. All experimental data, all analyses and all interpretations were generated and verified by the authors, who take full responsibility for the content.

## Data availability

The ENDOSPIN-29 dataset and the analysis code that reproduces every number and figure in this article are provided as Supplementary Information.

## Acknowledgements

Scanning electron microscopy was performed at UNAM, Bilkent University, and energy dispersive X-ray spectroscopy at Ordu University.

## Author contributions

N.B. designed and performed the electrospinning experiments, carried out the morphological characterization, curated the dataset, performed the computational analysis and wrote the manuscript. B.U. designed and performed the antibacterial, cytotoxicity and intracanal removal studies and contributed the calcium hydroxide formulation data. D.Ç.S. supervised the work, secured resources and revised the manuscript. All authors reviewed and approved the final manuscript.

